# Heterologous naringenin production in the filamentous fungus *Penicillium rubens*

**DOI:** 10.1101/2023.09.18.558247

**Authors:** Bo Peng, Lin Dai, Riccardo Iacovelli, Arnold J. M. Driessen, Kristina Haslinger

## Abstract

Naringenin is a natural product with several reported bioactivities and is the key intermediate for the entire class of plant flavonoids. The translation of flavonoids into modern medicine as pure compounds is often hampered by their low abundance in nature and difficult chemical synthesis. Here, we investigated new avenues toward producing high levels of naringenin in microbial hosts. *Penicillium rubens* is a well characterized and highly engineered traditional “workhorse” for the production of β-lactam antibiotics and cholesterol-lowering statins. We explored a secondary metabolite deficient *P. rubens* strain, *P. rubens* 4xKO, that was derived from an earlier industrial production strain as a promising microbial host for a recombinant flavonoid pathway. By integrating two plant genes encoding for enzymes in the naringenin biosynthesis pathway into the genome of this strain, we achieved a high naringenin titer in flask fermentations 36 h after feeding the precursor *p*-coumaric acid. Along with the rapid product accumulation of up to an 88% molar yield, we also observed rapid degradation of naringenin. Based on high-resolution mass spectrometric analysis, we identified the degradation products and proposed a naringenin degradation pathway in *P. rubens* 4xKO, which is distinct from other flavonoid-converting pathways reported in fungi. Our approach combines fundamental research with application-oriented microbial engineering, and our findings will pave the way to the more sustainable and economically feasible production of flavonoids for pharmaceutical and nutraceutical applications.

## 1. Introduction

Flavonoids are natural products found in various fruits, vegetables, and flowers. They belong to a class of plant secondary metabolites with a polyphenolic structure (Fig. 1). Plants use flavonoids for the growth and development of seedlings, the production of color and aromas to attract pollinators, and to protect themselves against different biotic and abiotic stresses^1,2^. For humans, flavonoids are an integral part of our diet and are mostly responsible for the color, taste, prevention of fat oxidation, and protection of vitamins and enzymes in food^3,4^. Additionally, flavonoids are reported to have several benefits on human health. This has been attributed to their antioxidant, antitumor, antiviral, anti-inflammatory, and neuroprotective activities, which have been reported in experiments with mammalian cell cultures^5–7^. These health-promoting effects make flavonoids highly attractive for nutraceutical, pharmaceutical, and cosmetic applications^2^.

**Figure 1:**
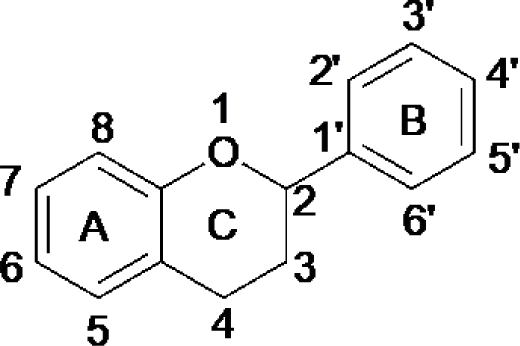
Basic structure of flavonoids.

Unfortunately, the current manufacturing routes do not provide scalable processes for large-scale production of pure flavonoids, hampering the study of these molecules for wider applications. The various extraction and purification steps needed for isolation from plant material, or the chemical synthesis come at high production costs and negatively impact the environment^8–10^. This is due to the low relative abundance of flavonoids in the plant tissues and the difficulty in separating them on a preparative scale since they exist as complex mixtures of structurally similar compounds^11^. Furthermore, the cultivation of plants for flavonoid extraction is rather inefficient because of long growing seasons^12,13^.

Therefore, the fermentative production of specific flavonoids, such as naringenin, an important precursor for many flavonoids, has attracted significant attention over the last 15 years. Several studies report the successful production of flavonoids using microbial hosts such as *Escherichia coli, Saccharomyces cerevisiae*, *Streptomyces clavuligerus,* and *Yarrowia lipolytica*^8,11,14^. However, to date, the yields obtained in these hosts are still too low to be competitive on the market, and sometimes expensive additives, e.g., inducers of gene expression and inhibitors of fatty acid synthesis, are needed to maximize production yields^15^. The major bottleneck in microbial naringenin production appears to be the limitation of free malonyl-Coenzyme A (malonyl-CoA) that is available for secondary metabolism in the host strain. Malonyl-CoA is the main precursor for fatty acid biosynthesis, an essential process in primary metabolism, and its abundance is strictly regulated to avoid waste of cellular resources. The key enzyme of naringenin biosynthesis, chalcone synthase (CHS) (Fig. 2A), is a type III polyketide synthase and directly competes with fatty acid biosynthesis for malonyl-CoA, since this is one of its natural substrates. Therefore, it has been crucial to increase the malonyl-CoA pool by engineering its upstream pathway and by suppressing fatty acid synthesis in microbial flavonoid producers^15,16^.

**Figure 2.**
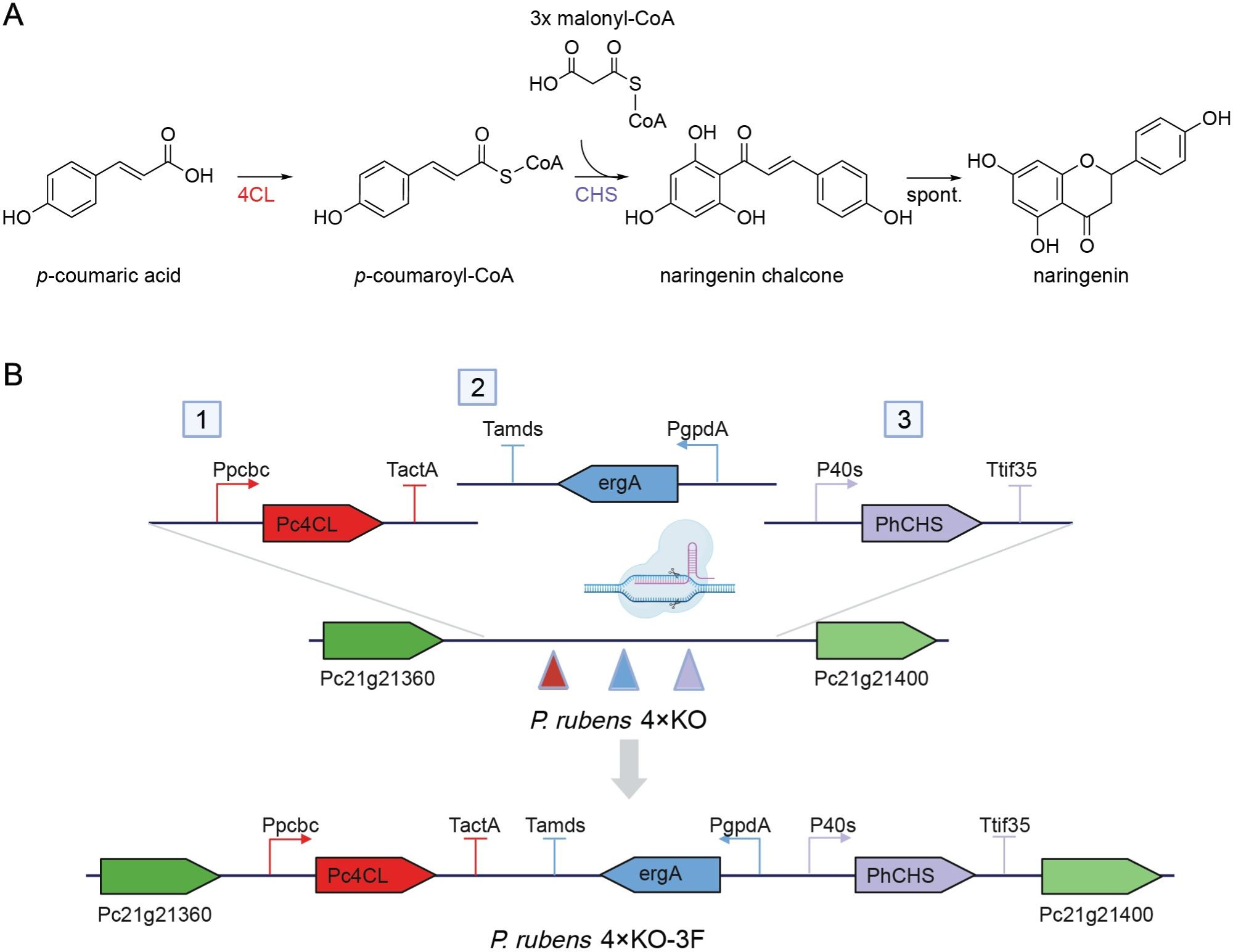
Biosynthesis pathway of naringenin and integration of the naringenin biosynthesis pathway into the *pen* locus of *P. rubens* 4xKO. (A) Biosynthesis pathway of naringenin. (4CL: 4-coumarate: CoA ligase, CHS: chalcone synthase). (B) Scheme for recombination of two (2 and 3) or three (1, 2, and 3) fragments (each fragment possesses homology arms of 100 bp) obtained by PCR into the intergenic region between Pc21g21360 and Pc21g21400. The obtained recombination strains were named *P. rubens* 4xKO-2F and *P. rubens* 4xKO-3F. Obtained clones were verified by colony PCR (Fig. S1).

The aforementioned bottleneck suggests that using a microbial host that is naturally gifted, or already engineered to produce high levels of secondary metabolites relying on malonyl-CoA, could be a good choice to recombinantly produce flavonoids. Therefore, we turned to derivatives of *P. rubens* Wisconsin 54–1255. These derivatives were previously engineered to produce β-lactam antibiotics but also shown to support the high-level production of cholesterol-lowering statins, which are malonyl-CoA dependent polyketides^17,18^. Specifically, we chose the secondary metabolite-deficient derivative of *P. rubens* DS68530, named *P. rubens* 4xKO. This strain was recently constructed via CRISPR/Cas9-based genome editing, via deletion of four highly expressed secondary metabolite gene clusters^19,20^. The low background of endogenous secondary metabolites simplifies the detection of target molecules, facilitates the downstream purification of the desired product and prevents that valuable resources are not utilized for unwanted secondary metabolites. This makes this strain an excellent option for the heterologous production of small molecules. To establish naringenin production in *P. rubens* 4xKO, we integrated the genes coding for CHS and 4-coumarate: CoA ligase (4CL) via CRISPR/Cas9-mediated engineering. After optimizing media composition and precursor feeding strategy, we achieved high molar yields of naringenin from the fed precursor *p*-coumaric acid. We also observed the ability of *P. rubens* to degrade naringenin and investigated the degradation pathway by metabolomics.

## 2. Materials & Methods

### 2.1 Strains, media, and culture conditions

*Escherichia coli* DH10β strain was used for cloning of transcription units. *P. rubens* 4xKO (Δ*penicillin*-BGC, Δ*chrysogine*-BGC, Δ*roquefortine*-BGC:: *amds*, Δ*hcpA*::*ble*, Δ*hdfA*) strain was used for the heterologous expression of the naringenin biosynthesis genes^20^. Spores of *P. rubens* stored on lyophilized rice grains were first germinated as precultures in YGG medium (in g/L): KCl, 8.0; glucose, 16.0; yeast nitrogen base (YNB), 6.66; citric acid, 1.5; K2HPO4, 6.0; and yeast extract, 2.0. Secondary metabolite-producing medium (SMP, pH 6.3) was prepared for secondary metabolite production, and the components of SMP medium are listed below (in g/L): K2HPO4, 2.12; KH2PO4, 5.1; CH3COONH4, 5.0; urea, 4.0; Na2SO4, 4.0; carbon source, 75.0; and supplemented with 4.0 ml/L Trace Element Solution^21^. Protoplasts were recovered for 5 to 7 days on solid transformation medium containing (in g/L): sucrose, 375.0; agar, 15.0; glucose, 10.0, and 4.0 ml/L Trace Element Solution, 27.0 ml/L Stock Solution A (KCl, 28.8; KH2PO4, 60.8; NaNO3, 240.0; at pH 5.5), 27.0 ml/L Stock Solution B (MgSO4·7H2O, 20.8), and pH adjusted around 7.0, when appropriate, supplemented with 1.1 μg/ml terbinafine hydrochloride (Sigma Aldrich, USA) for selection^21,22^. For sporulation, purification or preparation of lyophilized rice batches of *P. rubens* strains, R-agar medium was used, supplemented with 1.1 μg/ml terbinafine hydrochloride, and prepared as following (in g/L): agar 15.0; yeast extract, 5.0; MgSO4**·**7H2O, 0.05; NaCl, 18.0; CaSO4**·**2H2O, 0.25; KH2PO4, 0.06; CuSO4**·**5H2O, 0.01; NH4Fe(SO4)2**·**12H2O, 0.16; added with 7 mL 85% glycerol and 7.5 mL sugar beet molasse, provided by DSM-Firmenich (Delft, Netherlands). All *P. rubens* strains were incubated at 25 °C, and 200 rpm in 125 mL baffled flasks (Bellco, USA) for liquid medium.

### 2.2 Plasmid construction

All plasmids and primers used in this study are summarized in Tables 1 and S1. The genes of Pc4CL, 4-coumarate: CoA ligase from *Petroselinum crispum* (GenBank accession number KX671122.1), and PhCHS, chalcone synthase from *Petunia hybrida* (GenBank accession number KP284563.1) were ordered as synthetic genes from Integrated DNA Technologies (IDT, EU). All vectors were constructed via the Golden Gate technology-based Modular Cloning (MoClo) system using Type IIS restriction enzymes BpiI and BsaI as described previously^23^. To assemble *Pc4CL* and *PhCHS* into the MoClo entry vectors (level 0) pFL_0_1_Pc4CL and pFL_0_2_PhCHS, respectively, both ORF fragments were amplified with KAPA HiFi HotStart ReadyMix (Roche Diagnostics, Switzerland) with the primers (Table 1) that carried two BpiI restriction sites at the 5′-and 3′-end to allow cloning of the PCR products into vector pICH41308. The genes of interest, promoters, and terminators were then constructed into the MoClo transcription unit vectors (level 1) pICH47742 and pICH47761, with the BsaI restriction enzyme (Thermo Fisher Scientific, Waltham, MA)^24^. Both level 0 and level 1 plasmids were constructed to replace the *lacZ* gene of each backbone vector and then transformed into *E. coli* DH10β competent cells. Correctly assembled vectors were identified with blue-white screening, isolated by miniprep kit (Sigma Aldrich, USA) and analyzed by sequencing (Macrogen, Europe B.V.).

**Table 1.** Plasmids used in the study.

| Plasmid | Application | Template | Origin |
| --- | --- | --- | --- |
| pICH41308 | Level 0 cloning backbone for *pc4cl* and *phchs* | - | Addgene#47998<sup>23</sup> |
| pICH47742 | Level 1 cloning backbone for *pc4cl* | - | Addgene#48001<sup>23</sup> |
| pICH47761 | Level 1 cloning backbone for *phchs* | - | Addgene#48003<sup>23</sup> |
| pFTK013 | PpcbC Pc21g21380 promoter for *pc4cl* | *P. rubens* DS54468<sup>25</sup> | Addgene#171285<sup>24</sup> |
| pFTK081 | TactA ANIA_06542 *P. rubens* terminator for *pc4cl* | pDSM-JAK-108<sup>26</sup> | Addgene#171353<sup>24</sup> |
| pFTK012 | P40s AN0465 promoter for *phchs* | pDSM-JAK-108<sup>26</sup> | Addgene#171284<sup>24</sup> |
| pFTK076 | Ttif35 Pc22g19890 terminator for *phchs* | pDSM-JAK-108<sup>26</sup> | Addgene#171348<sup>24</sup> |
| pCP1_45 | Level 1 vector with *PgpdA-ergA-Tamds* as the terbinafine selection marker | *P. rubens* DS54468<sup>27</sup> | Pohl *et al.*, (2020)<sup>20</sup> |
| pFL_0_1_Pc4CL | Level 0 vector with *pc4cl* CDS | pETDuet:: *pc4cl*<sup>28</sup> | This study |
| pFL_0_2_PhCHS | Level 0 vector with *phchs* CDS | pETDuet:: *phchs*<sup>28</sup> | This study |
| pFL_1_1_Pc4CL | Level 1 vector with TU for donor DNA | - | This study |
| pFL_1_2_PhCHS | Level 1 vector with TU for donor DNA | - | This study |

CDS: coding sequence; TU: transcription unit.

Donor DNA fragments were amplified by PCR reactions from the level 1 plasmids, pFL_1_1_Pc4CL, pFL_1_2_PhCHS, and pCP1_45, introducing 100 bp long flanking regions on each end for homologous recombination. All three fragments were integrated into the *P. rubens* 4xKO chromosome at the original penicillin gene cluster chromosomal site (*pen* locus), 5’ of the *penDE* gene Pc21g21370 and 3’ of the *pcbAB* gene Pc21g21390 (Fig. 2B).

### 2.3 Fungal transformations

The preparation of protoplasts, transformation and colony screening were performed as described previously^20,22^. Approximately 2×10^7^ protoplasts, 8 μg donor DNA, and the pre-incubated mixture of 27 μg purified Cas9 protein and 4 μL synthesized single guide RNAs (sgRNAs) were mixed together for transformation. Cas9 protein was overexpressed in *E. coli* T7 Express lysY (New England Biolabs, UK) from pET28a/Cas9-Cys (Addgene plasmid # 53261), which was kindly provided by Hyongbum Kim via Addgene, and purified via Ni-NTA affinity chromatography^29^. The T7-sgRNA DNA templates were generated by PCR amplification with a pair of overlapping primers, and then the MEGAscript T7 Transcription Kit (Thermo Fisher Scientific, USA) was used to synthesize sgRNA. The transformed protoplasts were plated onto transformation media with terbinafine and incubated for 5-6 days at 25 °C with increased humidity for recovery^20,22^. Transformants were screened via colony PCR (Table S1, Fig. S1) using Phire Green Hot Start II PCR Master Mix (Thermo Fisher Scientific, USA) to confirm integration of donor DNA elements at the desired genomic locus and verified by sequencing. Correct transformants were grown on terbinafine containing R-agar plates for 5 to 6 days to produce spores and purified by three rounds of sporulation and genotype confirmation, to obtain genetically pure clones. For long-term storage, spores were inoculated on autoclaved-dried long-grain rice (Ben’s Original, USA), lyophilized, and stored at room temperature^20,21^.

### 2.4 Fermentation conditions, secondary metabolite extraction and analysis, and biomass measurement

The fermentation was performed in liquid culture; Three grains of rice with immobilized fungal spores (1.7 x 10^7^ spores/grain) were inoculated for 24 hours in 3 mL YGG medium before being transferred into 22 mL SMP medium in a 125 mL flask. After 1 - 4 days of cultivation, the precursor *p*-coumaric acid was added to the culture, and samples were taken periodically over the course of several days as described in the results section. To quantify naringenin concentration, 1 mL of culture was taken from the flask and mixed with 1 mL of methanol solution (100% methanol with 0.1% TFA). The mixture was centrifuged for 10 min at 13,000 rpm, and the supernatant was used for HPLC analysis. To identify secondary metabolites of *P. rubens* 4xKO variants, 2 mL samples were taken from the flask and extracted twice with 4 mL of ethyl acetate. The organic phase was collected and evaporated. After evaporation, 1 mL of 50% methanol (in water) was added to dissolve the crude extract and filtered by 0.45 μm PTFE syringe filter to remove insoluble particles prior to analysis. The samples were stored at −20 °C if not used immediately for high performance liquid chromatography-coupled mass spectrometry (HPLC-MS) analysis. For the biomass determination, mycelia were harvested by vacuum filtration over 0.45 μm cellulose filters (Sartorius, Germany) at indicated times. The collected biomass was dried at 60 °C for 60 h and then weighed.

### 2.5 Analysis and quantification of target compounds

Chemical standards for *p*-coumaric acid, naringenin, and phloretin were purchased from Sigma Aldrich (USA). Quantification of naringenin from the fermentation broths was achieved by high-performance liquid chromatography (HPLC, Shimadzu LC-10AT, equipped with a SPD-20A photodiode array detector) using a previously reported HPLC method^28^. Briefly, 10 μL of samples were injected into an Agilent Eclipse XDB-C18 (5 μm, 4.6×150 mm) column and separated with the following mobile phases: A: water + 0.1% trifluoroacetic acid (TFA); B: acetonitrile + 0.1% TFA. The following gradient was used: 15% B for 3 min, 15-90% B over 6 min; 90% B for 2 min; 90-15% B over 3 min, 15% B for 4 min; flow rate: 1 mL/min. *P*-coumaric acid and naringenin were identified by comparison to chemical standards. The peak areas were integrated and converted to concentrations based on calibration curves (Fig. S2) obtained with chemical standards.

The identity of secondary metabolites was assessed utilizing liquid chromatography-mass spectrometry (LC-MS) with a Waters Acquity Arc HPLC-MS system equipped with a 2998 PDA detector and a QDa single quadrupole mass detector. Samples (1 μL) were injected into and separated over an Xbridge BEH C18 (3.5 µm, 2.1×50 mm) column with the following mobile phases: A: water + 0.1% formic acid (FA); B: acetonitrile + 0.1% FA. The following gradient was used: 5% B for 2 min, 5-50% B over 15 min, 50-90% B over 4 min; 90% B for 3 min, 90-5% B over 6 min; flow rate: 0.25 mL/min. MS analysis was carried out in positive mode, with the following parameters: probe temperature 600 °C; capillary voltage 1.0 kV; cone voltage 15 V; scan range 100-1250 m/z.

High resolution tandem MS analyses were performed with a Shimadzu Nexera X2 HPLC system with binary LC20ADXR interfaced to a Q Exactive Plus Hybrid Quadrupole-Orbitrap Mass Spectrometer (Thermo Scientific). A 100×2.1 mm Kinetex EVO C18 reversed-phase column with 2.6 µm 100 Å particles (Phenomenex) was used for separation. The column and autosampler temperatures were set at 50 °C and 10 °C, respectively. The injection volume was 2 µL, and the flow was set at 0.25 mL/min. Mobile phases were used as same as HPLC-MS. The following gradient was used: 5% B for 2 min, 5-50% B over 32 min, 50-90% B over 8 min; 90% B for 3 min, 90-5% B over 5 min. MS and MS/MS analyses were performed with electrospray ionization in positive mode at a spray voltage of 3.5 kV, and sheath gas pressure of 60 psi, and auxiliary gas flow of 11 arbitrary units. The ion transfer tube temperature was 300 °C. Spectra were acquired in data-dependent mode with a survey scan at m/z 100-1650 at a resolution of 70,000 followed by MS/MS fragmentation of the top 5 precursor ions at a resolution of 17,500. A normalized collision energy (NCE) of 30 was used for fragmentation and fragmented precursor ions were dynamically excluded for 10 s.

## 3. Results

### 3.1 Validation of naringenin production in engineered *Penicillium rubens* 4xKO-2F and −3F

It is well established that the expression of a chalcone synthase (CHS) and a 4-coumarate: CoA ligase (4CL) in heterologous hosts in bacteria and yeast is sufficient to support the production of naringenin from fed *p*-coumaric acid. In particular, PhCHS from *Petunia hybrida* and Pc4CL from *Petroselinum crispum* are reported to be a highly efficient combination of enzymes^30–32^. Since *P. rubens* has been reported to express several native CoA ligases, with one of them even accepting *p*-coumaric acid as a substrate *in vitro*,^33^ we first tested if both plant enzymes need to be expressed in this new host to produce naringenin. We constructed the two strains *P. rubens* 4xKO-2F and −3F by genomic integration of the PhCHS encoding gene or the PhCHS and Pc4CL encoding genes, respectively, into the *pen* locus (Fig. 2a) and confirmed the integration by colony PCR (Fig. S1).

We then performed fermentations with the standard laboratory protocol in selective SMP medium (1.1 μg/mL terbinafine) with 75 g/L lactose as carbon source and with 1 mM *p*-coumaric acid as the precursor for naringenin production. When analyzing the culture extracts of the engineered variants by HPLC-MS, we observed a peak with the same retention time (RT) and mass-to-charge ratio (m/z) as the naringenin standard (Fig. 3). However, the titers in the *P. rubens* 4xKO-2F culture were extremely low. This suggests that one of the native CoA ligases can support the conversion of fed *p*-coumaric acid to coumaroyl-CoA, yet not efficiently. Co-expression of the two plant enzymes appears to be the better strategy to produce naringenin in *P. rubens*. In this co-expression strain (−3F), the highest titer of naringenin was about 0.05 mM 24 h after precursor addition. From 24 hours to 36 hours, the naringenin concentration decreased. This indicates that naringenin might also be rapidly degraded by the growing fungal cultures.

**Figure 3.**
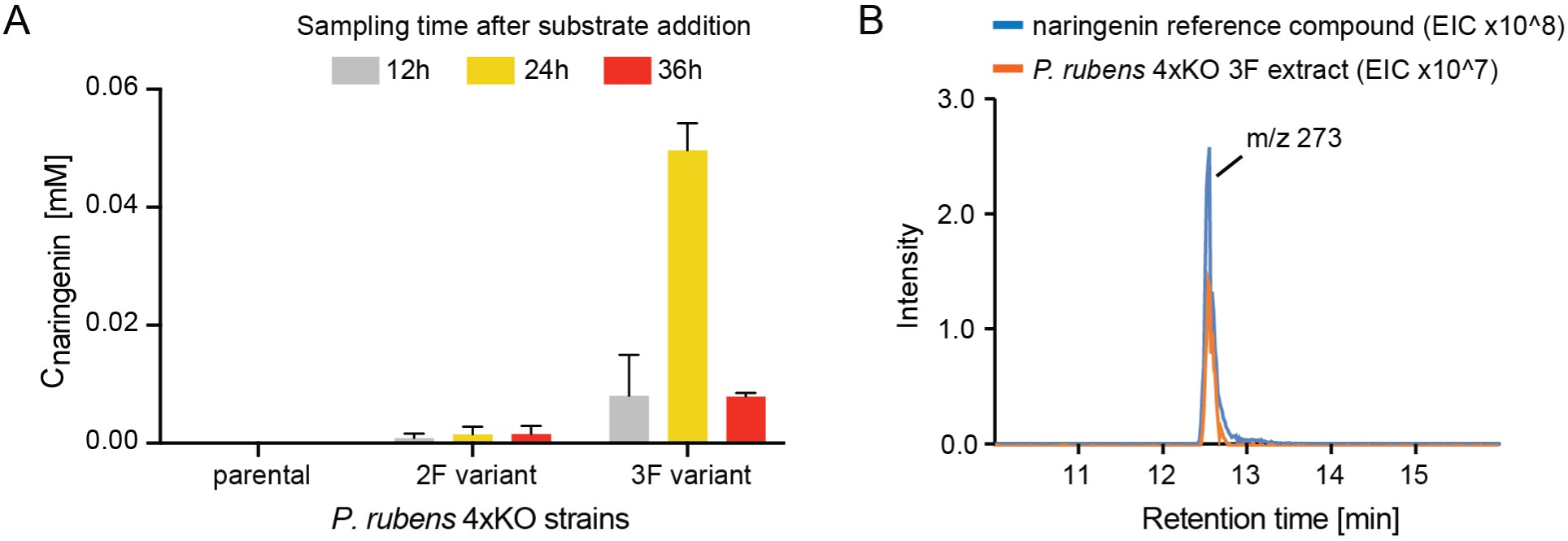
Comparison of *P. rubens* 4xKO variants towards naringenin production and chromatography of naringenin standard and samples. A) Naringenin titers detected in culture extracts of *P. rubens* 4xKO strains taken at different time points after precursor feeding. Data points represent mean +/-SD of biological replicates, n=3. B) Extracted ion chromatograms of the culture extract of *P. rubens* 4xKO-3F (orange) and the naringenin reference compound (blue) ([M+H]^+^ at m/z 273, RT=12.57 min).

### 3.2 Optimization of precursor feeding time

Based on the results shown above, we assumed that *P. rubens* degraded naringenin from 24 to 36 hours after precursor addition, when the fungal biomass was still increasing (data not shown). Thus, we set out to explore how precursor feeding time influences naringenin production. We varied the precursor feeding timepoint to 0, 24, 48, 72, or 96 h after inoculation of the precultures in the SMP medium, while keeping the other fermentation parameters the same. We took samples of the cultures every 24 h for several days and analyzed the extracts by LC-MS. The results show that the highest final naringenin titer was obtained in the cultures when the precursor was fed after 24 h of cultivation in the SMP medium at the 24 h sampling time point (Fig. 4A, around 0.05 mM naringenin). Surprisingly, we also detected another dominant compound in the cultures that were fed with the precursor after 72 h and 96 h of cultivation (Fig. 4B). Based on the m/z ratio of 275.0929 ([M + H]^+^) in tandem high resolution MS (Fig. S3), we predicted an elemental composition of C15H14O5 and hypothesized that this compound is phloretin, a reduced derivative of naringenin chalcone. We confirmed this hypothesis by comparing the retention time and m/z value to the commercially available reference compound. Upon closer inspection of all sample chromatograms, we also noticed that no phloretin can be detected in the samples in the other feeding conditions. One possible explanation for this phenomenon could be that in the absence of a chalcone isomerase, the naringenin chalcone generated by *P. rubens* 4xKO-3F cannot be converted into naringenin immediately and is then reduced to phloretin by native reductases. Thus, we concluded that feeding of *p*-coumaric acid after 24 h of cultivation in SMP medium followed by harvesting the culture the next day was the best strategy.

**Figure 4.**
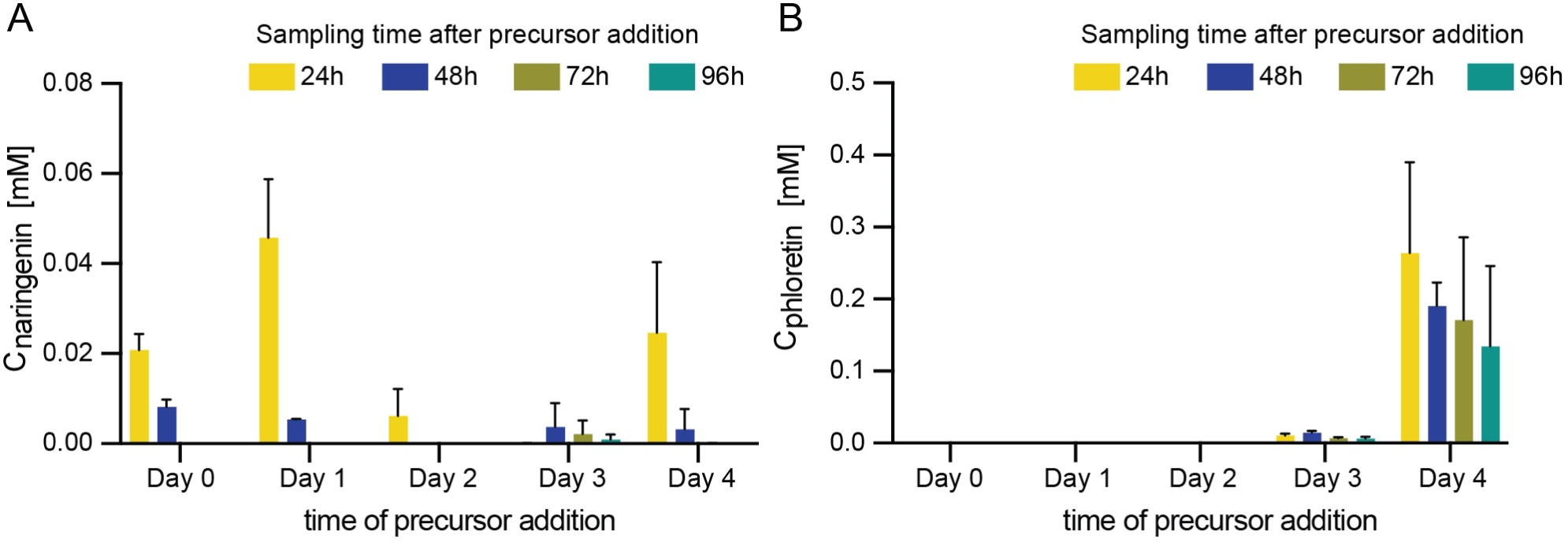
Time course of secondary metabolite accumulation in *P. rubens* 4xKO-3F with different precursor-feeding strategies. Fermentations were performed in SMP medium (lactose as a carbon source, pH 6.3) with 1 mM *p*-coumaric acid fed after 0, 24, 48, 72, and 96 h of cultivation in SMP medium. Samples were taken every 24 hours after feeding precursor, then extracted and analyzed by LC-MS. Data points represent mean +/-SD of biological triplicates, n=3. A) Naringenin titers (target compound). B) Phloretin titers (side product).

### 3.3 Optimization of naringenin production

Since the maximum titers of naringenin were very low with only 5% of the fed *p*-coumaric acid converted into the target product, we then set out to further improve the fermentation conditions. It is generally accepted that the media composition and pH can change the secondary metabolite profiles in filamentous fungi, and we therefore set out to explore different growth media^34,35^. In a quick pre-screening of carbon sources, we saw that utilizing glucose rather than lactose notably increased the final titers of naringenin (data not shown). So, for the pH optimization, the fermentations were carried out in SMP media with glucose as a carbon source and the pH of the phosphate buffer ranging from 6.3 to 8.0. We fed 1 mM *p*-coumaric acid after 24 h of cultivation in the modified SMP medium and took samples of the cultures every 12 h for HPLC analysis. The highest titers achieved with the different media were 0.3 mM (pH 6.3, 36 h after precursor feeding), 0.3 mM (pH 7.0, 24 h after precursor feeding), 0.2 mM (pH 7.5, 36 h after precursor feeding), and 0.55 mM (pH 8.0, 36 h after precursor feeding) (Fig. 5A), indicating that pH 8.0 gives the best result. Two days after precursor feeding, no naringenin was detected in any of the cultures, again demonstrating the subsequent degradation of the product.

**Figure 5.**
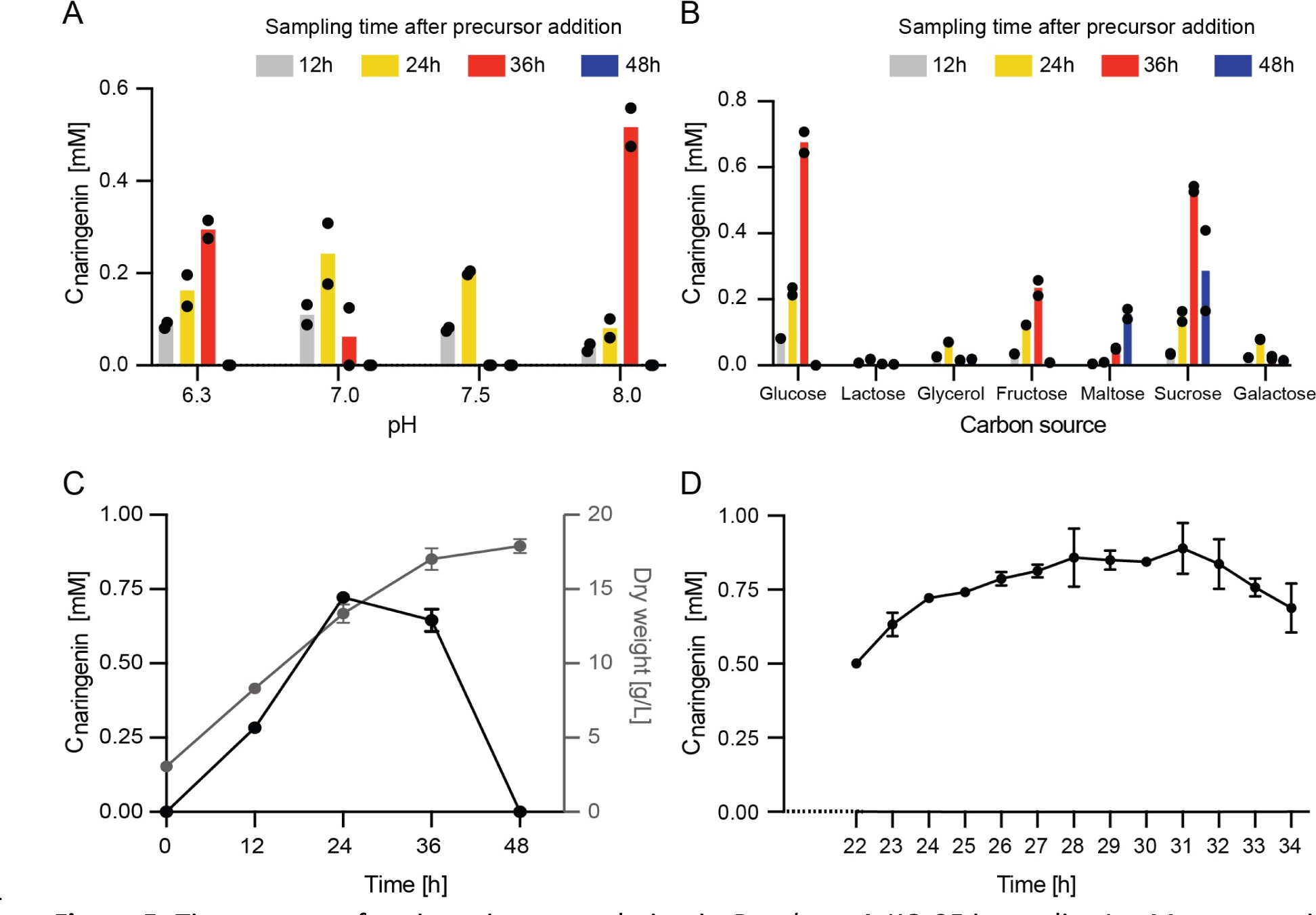
Time course of naringenin accumulation in *P. rubens* 4xKO-3F in media. 1 mM *p*-coumaric acid was fed after 24 h of cultivation. A) Naringenin titers obtained from cultures with different medium pH and glucose as a carbon source (n=2). B) Naringenin titers obtained from cultures with different carbon sources and a medium pH of 8.0 (n=2). C) Naringenin titers obtained from cultures in optimized media and *P. rubens* 4xKO-3F biomass accumulation during naringenin production sampled every 12 h (n=3) D) Naringenin titers obtained from cultures in optimized media sampled every hour from 22 h to 34 h after precursor addition (n=3). The highest titer of naringenin reached 0.88 mM from 1 mM *p*-coumaric acid.

After confirming the best pH condition, we repeated the screening for the best carbon source of the fermentation media and examined naringenin production under fermentation with different carbon sources (glucose, lactose, glycine, fructose, maltose, sucrose, and galactose) in the SMP medium (pH 8.0) (Fig. 5B). Consistent with our previous result, the naringenin titer produced in lactose-containing media is still low while glucose-containing media gives the highest titer. Cultivation in fructose- or sucrose-containing media, also supports higher conversion of fed *p*-coumaric acid and accumulation of naringenin than lactose. Thus, changing the carbon source and the pH of the media gave a boost in naringenin titer at the 36 h time point by one order of magnitude. From 36 h to 48 h, the concentration of naringenin in the cultures decreased.

To further improve naringenin titers, we next performed another time course experiment to optimize the time point for harvesting the culture and characterize the growth phenotype of the *P. rubens* 4xKO-3F strain. Since the antifungal terbinafine was still used in all fermentations described thus far, we noticed poor growth in most experiments and decided to proceed without this additive. The genes encoding PhCHS and Pc4CL are integrated into the genome and the resulting strain was thoroughly purified. Therefore, this additional selection is not necessary. We cultivated the engineered strain under the optimized conditions (modified SMP medium with pH 8.0 and glucose as carbon source), fed the precursor 1 mM *p*-coumaric acid after 24 h, and took samples at a 12 h interval, with additional sampling every hour between 22 and 34 h (Fig. 5C and 5D). In the first 24 h, naringenin accumulated in the fermentation broth (from 0 to 0.7 mM), and the biomass increased linearly from 3.0 g/L to 13.3 g/L. Then from 24 to 31 h, the titer of naringenin reached a peak (around 0.88 mM naringenin), and the biomass accumulation slowed down (from 13.3 g/L to 17.0 g/L in 12 hours). After 31 h of cultivation, the concentration of naringenin began to decrease and the culture reached the stationary phase. At the 48 h time point, no naringenin was detected in the culture. The highest titer and molar yield (0.88 mM and 88%, respectively) were much higher than in our previous experiments and indicate that *P. rubens* can be pushed to produce high amounts of naringenin with an almost stochiometric conversion of *p*-coumaric acid into naringenin.

### 3.4 Naringenin degradation in *P. rubens* 4xKO

Since fungi play an important role in natural ecosystems for degrading biomass, it is well known that they have catabolic pathways to degrade and utilize plant polymers and small organic compounds. In the literature, several filamentous fungi and yeasts have been described to modify flavonoids by oxidative processes or glycosylation^36–39^, yet complete degradation of the scaffold has only been reported in bacteria^40,41^. For example, Marin *et al.* investigated naringenin degradation in the β-proteobacterium *Herbaspirillum seropedicae* SmR1^40,41^. They pinpointed a gene cluster responsible for the degradation and identified key intermediates by high resolution tandem MS and nuclear magnetic resonance. While Braune *et al* ^42^. demonstrated that an oxygen-sensitive NADH-dependent reductase from *Eubacterium ramulus* could cleave naringenin, eriodictyol, liquiritigenin, and homoeriodictyol.

With these studies in mind, we performed a time course experiment to investigate whether any of the heterologously expressed plant enzymes were involved in naringenin degradation and to investigate the intermediates of naringenin degradation. We cultured *P. rubens* 4xKO and *P. rubens* 4xKO-3F for one day in modified SMP medium as described above (pH 8.0, glucose as carbon source) before adding 1 mM naringenin. Next, samples were collected every 24 h for HPLC and HPLC-MS analysis with an additional sample taken after 6 h. For both strains, we observed that the concentration of naringenin decreases linearly over 72 h until most naringenin was degraded (final concentration 0.06 mM in cultures of the parental strain and 0.037 mM in the 3F strain) (Fig. 6A). Furthermore, the biomass increases at a steady rate even after 48 h to a higher amount of cell dry weight that in all previous experiments (23.5 g/L for parental strain, 26 g/L for 3F) (Fig. 6B). Since the parental strain degrades naringenin, it can be concluded that the heterologously expressed plant enzymes are not required to degrade naringenin.

**Figure 6.**
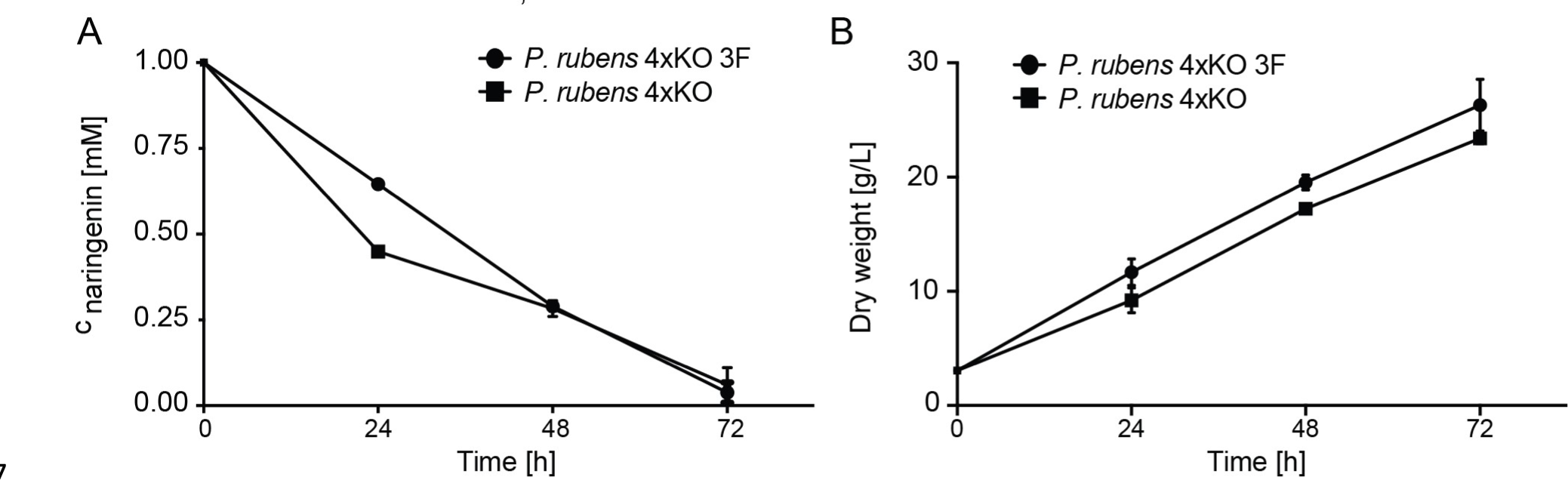
Time course of naringenin degradation in *P. rubens* 4xKO and *P. rubens* 4xKO-3F. Naringenin acid (1 mM) was fed after 24 h of cultivation in optimized SMP media. Data points represent mean +/-SD of biological replicates, n=3. A) Naringenin titer over time. B) Biomass accumulation over time.

Next, we compared the peak areas, retention times and m/z values of new peaks in the low-resolution HPLC-MS chromatograms of the various samples from both strains and found that there were also no variations in the degradation products of naringenin between *P. rubens* 4xKO and *P. rubens* 4xKO-3F. Thus, we performed high resolution tandem MS on the samples taken from the *P. rubens* 4xKO and *P. rubens* 4xKO-3F cultures to attempt to identify the major compounds (Table 2, Figure 7). At time point 0 h, we only detected naringenin in the culture, but after 6 h of cultivation we observed compound 2 with an m/z of 342.1364 ([M+H]^+^). Compound 2 persisted in the culture media for 72 hours, suggesting that it could be a dead-end product in *P. rubens* 4xKO. After 24 h of cultivation, compounds 3a, 3b, 4a, 4b, 4c, 5a, and 5b were found in the culture. Further analysis revealed that compounds 3a and 3b are isomers with an identical predicted molecular formula (C15H12O6) at m/z 289.0728 ([M+H]^+^), and were detected at RT 9.89 and 10.55 min, respectively. Based on the fragmentation pattern in MS^2^ with two characteristic fragments (m/z 169.0147 and 147.0453), these two features likely correspond to carthamidin and isocarthamidin with an additional hydroxyl group on the A-ring compared to naringenin^43–45^. Compounds 4a, 4b, and 4c are isomers with the predicted molecular formula (C11H10O4) at m/z 207.0669 ([M+H]^+^), detected at RT 1.61, 4.56, and 7.47 min. Compounds 5a and 5b are isomers with the predicted molecular formula (C10H10O2) at m/z 163.0768 ([M+H]^+^) detected at RT 2.62 and 9.12 min. After 72 h, compounds 4a-4c were degraded, but compounds 2, 5a, and 5b remained in the culture media. Based on the m/z and the fragmentation patterns in MS^2^, we assume that compounds 3a-5b are the same intermediates that were described for the degradation of naringenin in *H. seropedicae* SmR1^41^. In *H. seropedicae* SmR1, naringenin degradation begins from hydroxylation of the A-ring on c8 (isocarthamidin), followed by the loss of the A-ring putatively as oxaloacetate to form the intermediate 5-(3,4-dihydroxyphenyl)-3-oxovalero-δ-lactone. This lactone is then hydroxylated in the former c2 position of the former B-C ring, the C-ring opens, is decarboxylated and dehydrated to form the final product (*3E*)-4-(4-hydroxyphenyl) but-3-en-3one^41^. In the cultures of *P. rubens* 4xKO, we detected the expected m/z values for the hydroxylated flavonoid (m/z 289.0728 [M+H]^+^, compounds 3a/b), the lactone intermediate (m/z 207.0503 [M+H]^+^, compounds 4a-c) and the final product (m/z 163.0768 [M+H]^+^, compounds 5a/b) but not for the other intermediates. This suggests that flavonoid degradation in *P. rubens* 4xKO follows a similar pathway as described for *H. seropedicae* SmR1 (Fig. 8). The oxaloacetate that is released in this proposed pathway may feed into primary metabolism and thus boost biomass formation.

**Figure 7.**
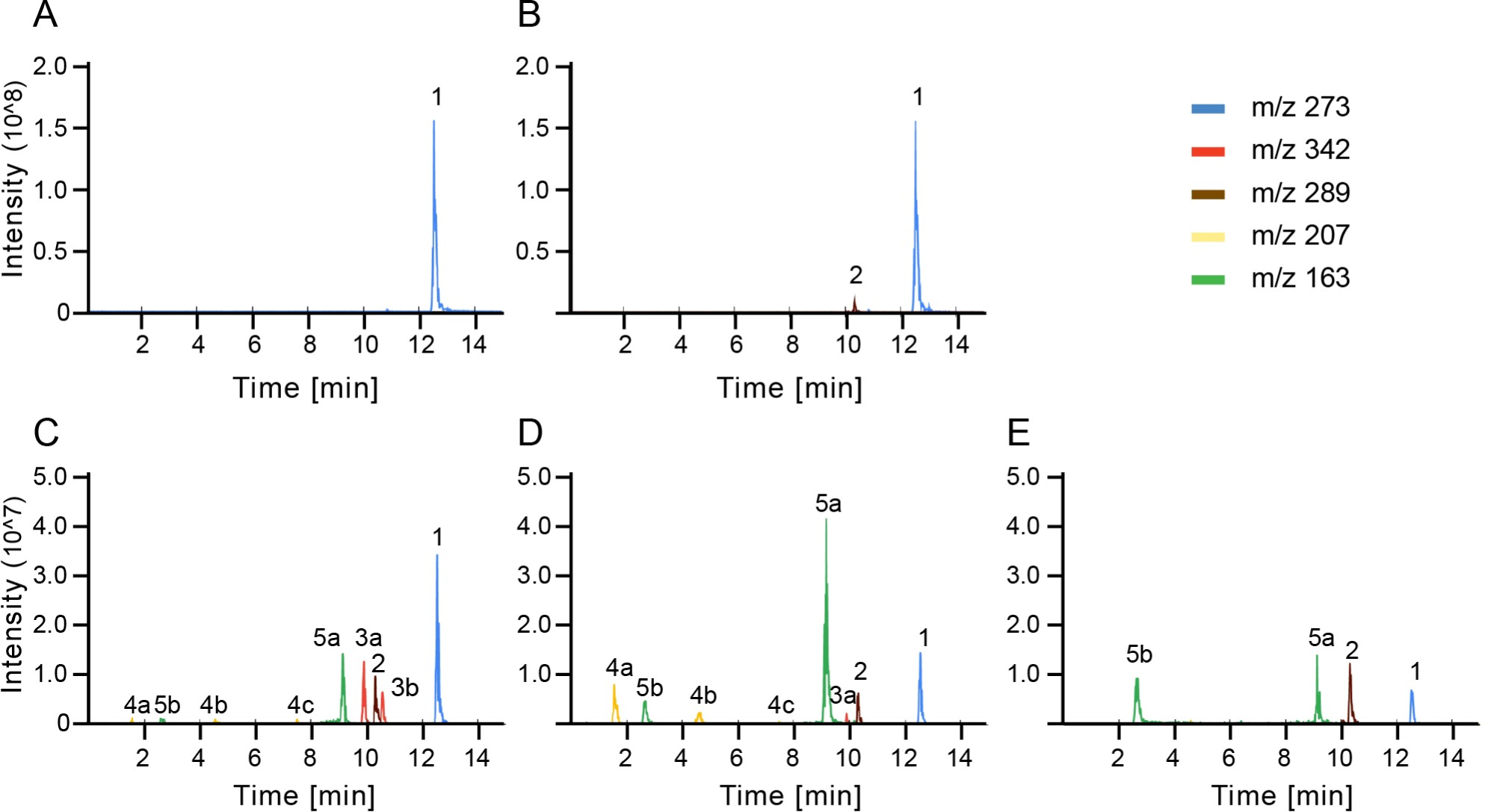
Extracted ion chromatograms of the major intermediates from naringenin degradation by *P. rubens* 4xKO. A) 0 h, B) 6 h, C) 24 h, D) 48 h, E) 72 h time point. The strain was grown at 25 °C in modified SMP medium containing 1 mM naringenin.

**Figure 8.**
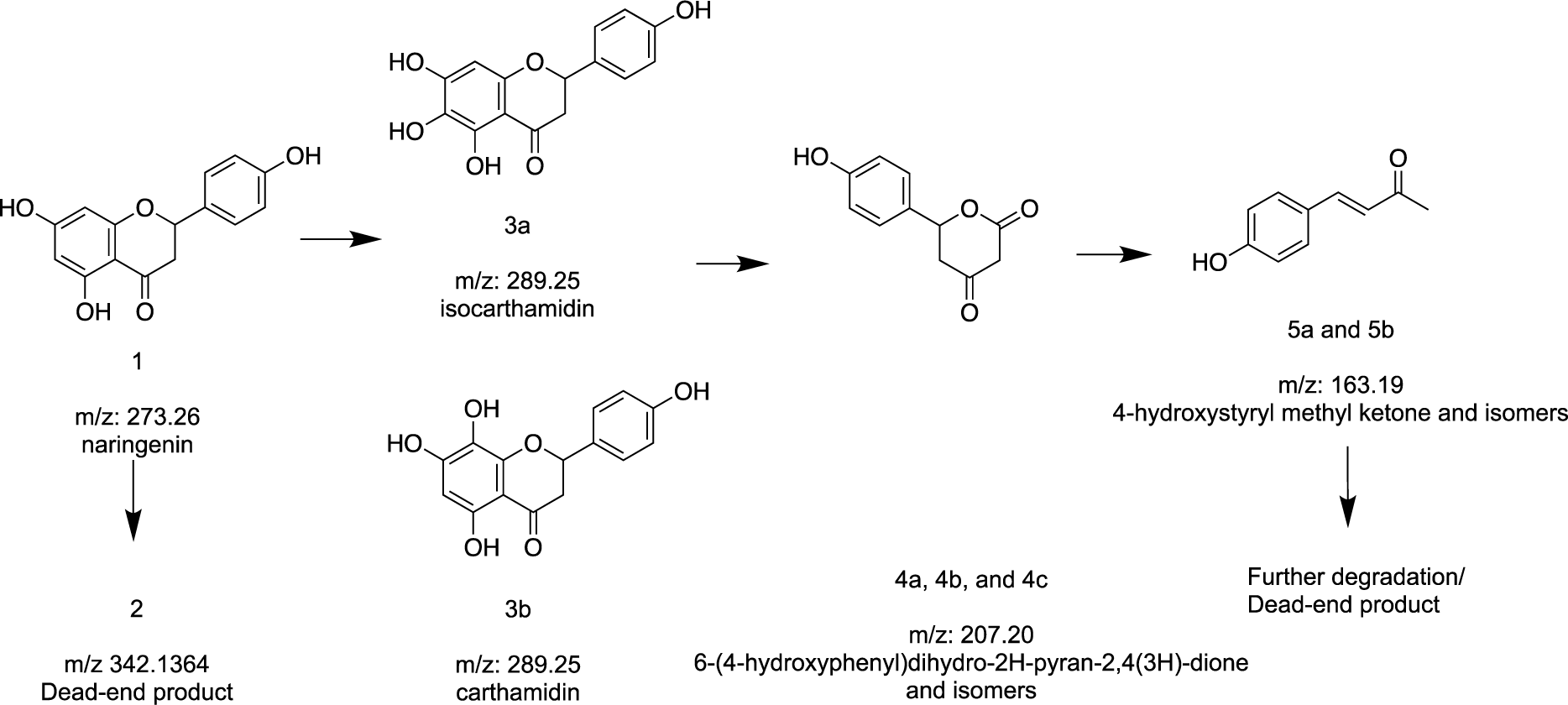
Proposed naringenin degradation pathway in *P. rubens* 4xKO.

**Table 2.**
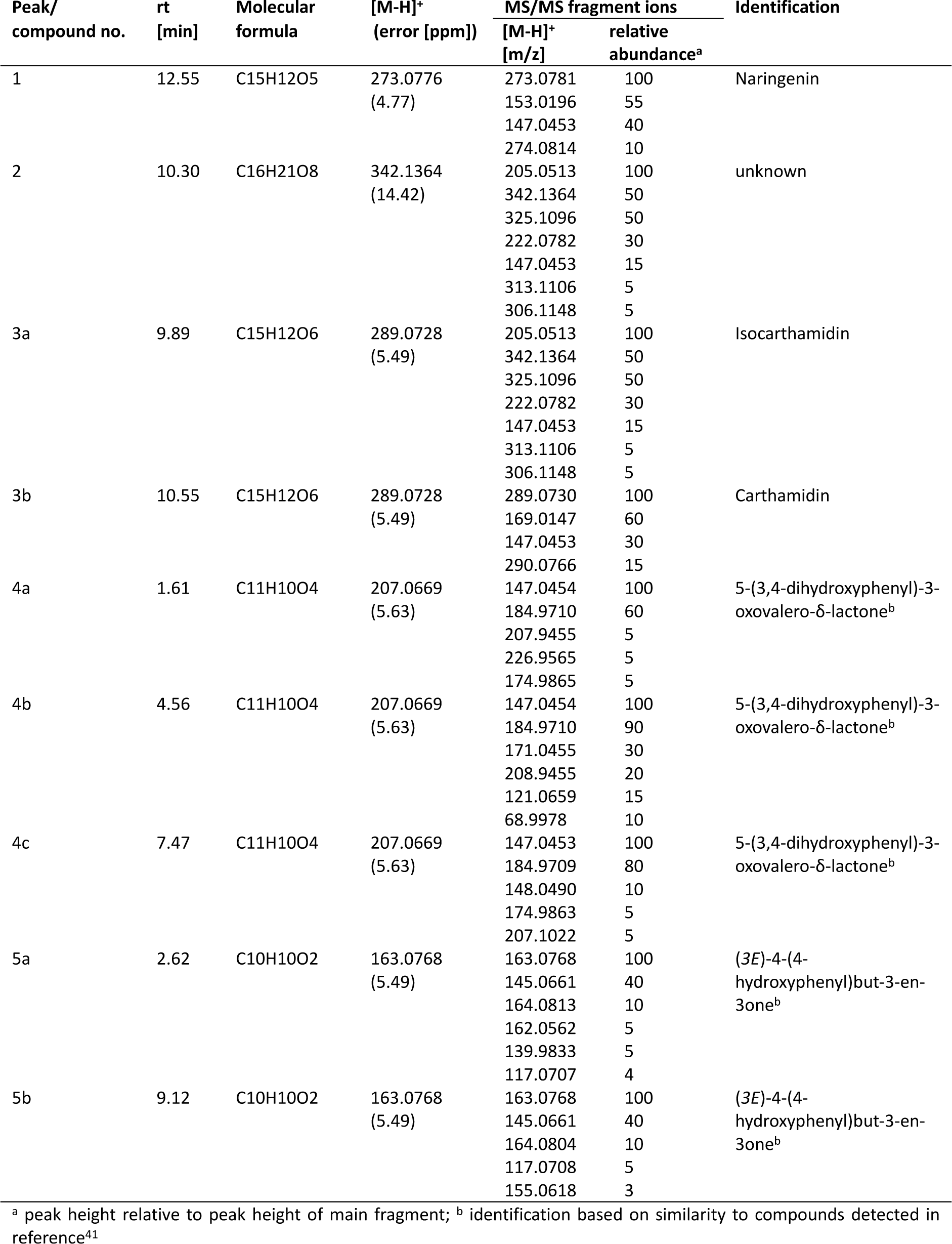
Characteristics of the nine major peaks detected by high resolution tandem MS in the extracts <u>of *P. rubens* 4xKO cultured in the presence of 1 mM naringenin.</u>.

Since Marin et al.^40,41^ also identified a gene cluster that is responsible for naringenin degradation in *H. seropedicae* SmR1, we looked for homologues of these genes in the published *P. rubens* Wisconsin 54-1255 genome (GCA_000226395)^46^. Although we were unable to identify a syntenic gene cluster, we found several monooxygenases and cupin-domain enzymes that could catalyze these reactions in this degradation pathway. It is of course also possible that filamentous fungi use different enzymes to achieve the same result.

## 4. Discussion and conclusions

In this study, we set out to assess the potential of *P. rubens* derivative strains to produce the polyketide naringenin, which is an important intermediate in the flavonoid metabolic network and a natural product with various health benefits^2,47–49^. By integrating only two plant genes encoding for two flavonoid pathway enzymes into the *P. rubens* 4xKO genome, we were able to detect low levels of naringenin produced from a fed precursor. After optimization of the fermentation conditions (precursor feeding time, medium pH, and carbon source,) we achieved a high final titer of naringenin (0.88 mM), which corresponds to 88% molar yield from the fed precursor *p*-coumaric acid in flask fermentation. We observed that changing the carbon source from lactose, which is optimal for penicillin production, to glucose led to an increase in naringenin titer by one order of magnitude. Changing the pH of the medium from 6.3 to 8.0 led to a further increase by about 1.5-fold. The media composition, especially the carbon source, affects the growth rate of *P. rubens* and since the heterologous pathway genes in the 3F strain are constitutively expressed, this has a direct impact on the availability of the biocatalysts. It is also possible that the modified medium slows down competing pathways, for example by negatively affecting the expression of native monooxygenases and reductases that facilitate the degradation of the newly built product. In fact, we observed rapid degradation of the newly formed naringenin throughout our experiments. We confirmed that also the parental strain is capable of degrading fed naringenin, which suggests that the heterologously expressed plant enzymes do not contribute to the degradation. When analysing the intermediates of naringenin degradation, we observed that they differ from known intermediates from fungal conversions of flavonoids reported before^50^. Instead, they are highly similar to the degradation products identified in *H. seropedicae* SmR1^41^. In this β-proteobacterium, it was demonstrated that an FAD-dependent monooxygenase, FdeE (Hsero_1007), performs the first catalytic step forming the 8-hydroxylated intermediate, and that a dioxygenase, FdeC (Hsero_1005), is involved in cleaving the A-ring ^40,41^. Several other enzymes encoded in the same gene cluster were further implicated in the degradation pathway, yet no experimental verification has been reported^41^. Our efforts to identify a homologous gene cluster in the *P. rubens* Wisconsin 54–1255 genome has not returned any hits. Therefore, a deeper investigation of the homologues of the individual enzymes is necessary to pinpoint the responsible genes in *P. rubens*. Once these are identified, it may be possible to eliminate this competing pathway and thereby engineer *P. rubens* into an even more powerful cell factory for flavonoids. Even in this unoptimized *P. rubens* strain, we have achieved final titers and molar yields for naringenin that are competitive compared to previous reports in other microbial hosts^15,32,51–55^.

In our initial experiments before optimizing the timing of precursor feeding and media composition, we also observed phloretin as a byproduct. Phloretin is a chemical widely applied in medical and cosmetic industries due to its skin-lightening property^38,56^. The concentration of phloretin reached approximately 0.25 mM from 1 mM *p*-coumaric acid, highlighting again the potential of the engineered *P. rubens* 4xKO-3F for the production of chalcones and flavonoids without further manipulation of the intrinsic malonyl-CoA pool. By integrating additional genes encoding for upstream or downstream enzymes, it will be possible to extend this minimal pathway. For instance, integrating a gene for tyrosine ammonia lyase would allow *de novo* synthesis of naringenin from *P. rubens* primary metabolites. Expanding the pathway with flavone synthase would give access to flavones from naringenin, while other oxidoreductases would lead to other types of flavonoids such as flavonols or anthocyanins^54,57^.

## Abbreviations and nomenclature

4CL: 4-coumarate: CoA ligase
CHS: chalcone synthase
CRISPR: Clustered Regularly Interspaced Short Palindromic Repeats
EIC: Extracted ion chromatogram
HR: homologous recombination
HPLC: High Performance Liquid Chromatography m/z, mass-to-charge ratio
MS: mass spectrometry RT, retention time
TFA: trifluoroacetic acid

## Supporting Information

Gene and primer sequences, evidence for genomic integration, HPLC calibration curves and high resolution tandem MS analysis for phloretin, carthamidin and isocarthamidin (PDF).

## Acknowledgment

We thank Dr. László Mózsik for his help in the initial design of the genome editing strategy and Prof. Roel Bovenberg from DSM-Fermenich, Delft, for support and valuable discussions. We thank the interfaculty mass spectrometry center of the University of Groningen and the University Medical Center Groningen for high resolution tandem MS analysis. Figures were prepared with elements from Biorender.com.

## Author contributions

KH, AJMD, and BP conceived the study with contributions from LD and RI; BP and LD cloned plasmids, prepared transformation, and performed fermentations under the guidance of RI; BP, LD, and KH analyzed biochemical data; KH and BP wrote the manuscript with contributions from LD, RI, and A.J.M.D; all authors have read and approved the final version of the manuscript.

## For table of contents only

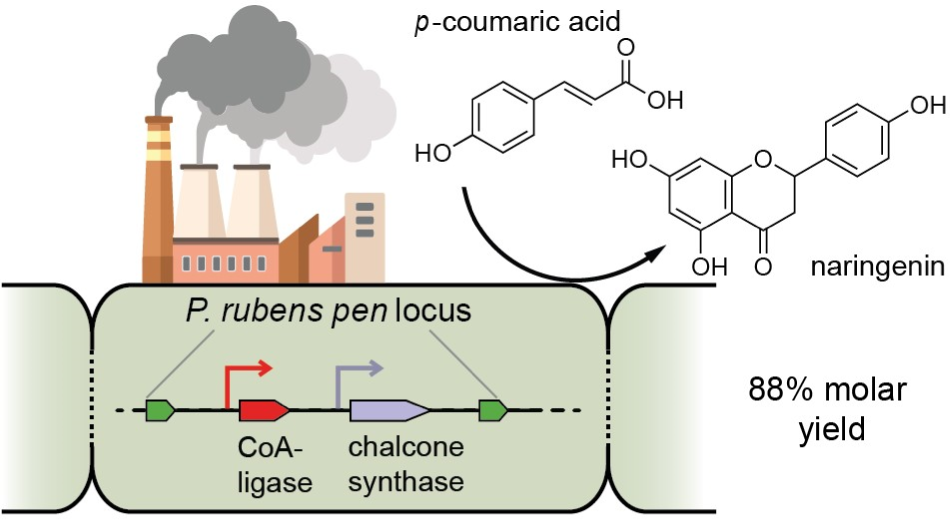

## References

1. Griesbach RJ. Biochemistry and genetics of flower color. Plant Breed Rev. 2010;25:89–114.

2. Panche AN, Diwan AD, Chandra SR. Flavonoids: An overview. J Nutr Sci. 2016;5. doi:10.1017/jns.2016.41

3. Huvaere K, Skibsted LH. Flavonoids protecting food and beverages against light. J Sci Food Agric. 2015;95(1):20–35.

4. Yao LH, Jiang YM, Shi J, et al. Flavonoids in food and their health benefits. Plant Foods for Human Nutrition. 2004;59(3):113–122. doi:10.1007/s11130-004-0049-7

5. Rozmer Z, Perjési P. Naturally occurring chalcones and their biological activities. Phytochemistry Reviews. 2016;15(1):87–120. doi:10.1007/s11101-014-9387-8

6. Roohbakhsh A, Parhiz H, Soltani F, Rezaee R, Iranshahi M. Molecular mechanisms behind the biological effects of hesperidin and hesperetin for the prevention of cancer and cardiovascular diseases. Life Sci. 2015;124:64–74. doi:10.1016/j.lfs.2014.12.030

7. Dong W, Wei X, Zhang F, et al. A dual character of flavonoids in influenza A virus replication and spread through modulating cell-autonomous immunity by MAPK signaling pathways. Sci Rep. 2014;4(1):7237. doi:10.1038/srep07237

8. Marsafari M, Samizadeh H, Rabiei B, Mehrabi A, Koffas M, Xu P. Biotechnological Production of Flavonoids: An Update on Plant Metabolic Engineering, Microbial Host Selection, and Genetically Encoded Biosensors. Biotechnol J. 2020;15(8):e1900432. doi:10.1002/biot.201900432

9. Zhang W, Yi D, Gao K, et al. hydrolysis of Scutellarin and Related Glycosides to Scutellarein and the Corresponding Aglycones. J Chem Res. 2014;38(7):396–398. doi:10.3184/174751914X14017253941699

10. Morikawa T, Wang LB, Nakamura S, et al. Medicinal Flowers. XXVII. New flavanone and chalcone glycosides, arenariumosides I, II, III, and IV, and tumor necrosis factor-alpha inhibitors from everlasting flowers of *Helichrysum arenarium*. Chem Pharm Bull (Tokyo). 2009;57(4):361–367. doi:10.1248/cpb.57.361

11. Lv Y, Marsafari M, Koffas M, Zhou J, Xu P. Optimizing Oleaginous Yeast Cell Factories for Flavonoids and Hydroxylated Flavonoids Biosynthesis. ACS Synth Biol. 2019;8(11):2514–2523. doi:10.1021/acssynbio.9b00193

12. Wang Y, Chen S, Yu O. Metabolic engineering of flavonoids in plants and microorganisms. Appl Microbiol Biotechnol. 2011;91(4):949–956. doi:10.1007/s00253-011-3449-2

13. Wang H, Wang W, Li H, Zhang P, Zhan J, Huang W. Expression and tissue and subcellular localization of anthocyanidin synthase (ANS) in grapevine. Protoplasma. 2011;248(2):267–279. doi:10.1007/s00709-010-0160-6

14. Álvarez-Álvarez R, Botas A, Albillos SM, Rumbero A, Martín JF, Liras P. Molecular genetics of naringenin biosynthesis, a typical plant secondary metabolite produced by *Streptomyces clavuligerus*. Microb Cell Fact. 2015;14(1):178. doi:10.1186/s12934-015-0373-7

15. Xu P, Ranganathan S, Fowler ZL, Maranas CD, Koffas MAG. Genome-scale metabolic network modeling results in minimal interventions that cooperatively force carbon flux towards malonyl-CoA. Metab Eng. 2011;13(5):578–587. doi:10.1016/j.ymben.2011.06.008

16. Kim BG, Lee H, Ahn JH. Biosynthesis of Pinocembrin from Glucose Using Engineered *Escherichia coli*. J Microbiol Biotechnol. 2014;24(11):1536–1541. doi:10.4014/jmb.1406.06011

17. Weber SS, Bovenberg RAL, Driessen AJM. Biosynthetic concepts for the production of β-lactam antibiotics in *Penicillium chrysogenum*. Biotechnol J. 2012;7(2):225–236. doi:10.1002/biot.201100065

18. McLean KJ, Hans M, Meijrink B, et al. Single-step fermentative production of the cholesterol-lowering drug pravastatin via reprogramming of *Penicillium chrysogenum*. Proc Natl Acad Sci U S A. 2015;112(9):2847–2852. doi:10.1073/pnas.1419028112

19. Pohl C, Kiel JAKW, Driessen AJM, Bovenberg RAL, Nygård Y. CRISPR/Cas9 Based Genome Editing of *Penicillium chrysogenum*. ACS Synth Biol. 2016;5(7):754–764. doi:10.1021/acssynbio.6b00082

20. Pohl C, Polli F, Schütze T, et al. A *Penicillium rubens* platform strain for secondary metabolite production. Sci Rep. 2020;10(1):7630. doi:10.1038/s41598-020-64893-6

21. Kovalchuk A, Weber SS, Nijland JG, Bovenberg RAL, Driessen AJM. Fungal ABC transporter deletion and localization analysis. Methods in Molecular Biology. 2012;835:1–16. doi:10.1007/978-1-61779-501-5_1

22. Mózsik L, Hoekzema M, de Kok NAW, Bovenberg RAL, Nygård Y, Driessen AJM. CRISPR-based transcriptional activation tool for silent genes in filamentous fungi. Sci Rep. 2021;11(1):1118. doi:10.1038/s41598-020-80864-3

23. Weber E, Engler C, Gruetzner R, Werner S, Marillonnet S. A modular cloning system for standardized assembly of multigene constructs. Peccoud J, ed. PLoS One. 2011;6(2):e16765. doi:10.1371/journal.pone.0016765

24. Mózsik L, Pohl C, Meyer V, Bovenberg RAL, Nygård Y, Driessen AJM. Modular Synthetic Biology Toolkit for Filamentous Fungi. ACS Synth Biol. 2021;10(11):2850–2861. doi:10.1021/acssynbio.1c00260

25. Polli F, Meijrink B, Bovenberg RAL, Driessen AJM. New promoters for strain engineering of *Penicillium chrysogenum*. Fungal Genetics and Biology. 2016;89:62–71. doi:10.1016/j.fgb.2015.12.003

26. Mózsik L, Büttel Z, Bovenberg RAL, Driessen AJM, Nygård Y. Synthetic control devices for gene regulation in *Penicillium chrysogenum*. Microb Cell Fact. 2019;18(1):203. doi:10.1186/s12934-019-1253-3

27. Sigl C, Handler M, Sprenger G, Kürnsteiner H, Zadra I. A novel homologous dominant selection marker for genetic transformation of *Penicillium chrysogenum*: Overexpression of squalene epoxidase-encoding ergA. J Biotechnol. 2010;150(3):307–311. doi:10.1016/j.jbiotec.2010.09.941

28. Peng B, Zhang L, He S, et al. Engineering a plant polyketide synthase for the biosynthesis of methylated flavonoids. preprint on bioRxiv. Published online 2022:1–27. 10.1101/2022.10.02.510496

29. Ramakrishna S, Kwaku Dad AB, Beloor J, Gopalappa R, Lee SK, Kim H. Gene disruption by cell-penetrating peptide-mediated delivery of Cas9 protein and guide RNA. Genome Res. 2014;24(6):1020–1027. doi:10.1101/gr.171264.113

30. Santos CNS, Koffas M, Stephanopoulos G. Optimization of a heterologous pathway for the production of flavonoids from glucose. Metab Eng. 2011;13(4):392–400. doi:10.1016/j.ymben.2011.02.002

31. Li H, Gao S, Zhang S, Zeng W, Zhou J. Effects of metabolic pathway gene copy numbers on the biosynthesis of (2S)-naringenin in *Saccharomyces cerevisiae*. J Biotechnol. 2021;325:119–127. doi:10.1016/j.jbiotec.2020.11.009

32. Akram M, Rasool A, An T, Feng X, Li C. Metabolic engineering of *Yarrowia lipolytica* for liquiritigenin production. Chem Eng Sci. 2021;230:116177. doi:10.1016/j.ces.2020.116177

33. Koetsier MJ, Jekel PA, van den Berg MA, Bovenberg RAL, Janssen DB. Characterization of a phenylacetate–CoA ligase from *Penicillium chrysogenum*. Biochemical Journal. 2009;417(2):467–476. doi:10.1042/BJ20081257

34. Luo Z, Li Y, Mousa J, et al. *Bbmsn2* acts as a pH-dependent negative regulator of secondary metabolite production in the entomopathogenic fungus *Beauveria bassiana*. Environ Microbiol. 2015;17(4):1189–1202. doi:10.1111/1462-2920.12542

35. VanderMolen KM, Raja HA, El-Elimat T, Oberlies NH. Evaluation of culture media for the production of secondary metabolites in a natural products screening program. AMB Express. 2013;3(1):71. doi:10.1186/2191-0855-3-71

36. Álvarez-Álvarez R, Botas A, Albillos SM, Rumbero A, Martín JF, Liras P. Molecular genetics of naringenin biosynthesis, a typical plant secondary metabolite produced by Streptomyces clavuligerus. Microb Cell Fact. 2015;14(1):1–12. doi:10.1186/s12934-015-0373-7

37. Sordon S, Popłoński J, Huszcza E. Microbial glycosylation of flavonoids. Pol J Microbiol. 2016;65(2):137–151. doi:10.5604/17331331.1204473

38. Jiang C, Liu X, Chen X, et al. Raising the production of phloretin by alleviation of by-product of chalcone synthase in the engineered yeast. Sci China Life Sci. 2020;63(11):1734–1743. doi:10.1007/s11427-019-1634-8

39. Sordon S, Popłoński J, Tronina T, Huszcza E. Regioselective O-glycosylation of flavonoids by fungi *Beauveria bassiana*, *Absidia coerulea* and *Absidia glauca*. Bioorg Chem. 2019;93:102750. doi:10.1016/j.bioorg.2019.01.046

40. Marin AM, Souza EM, Pedrosa FO, et al. Naringenin degradation by the endophytic diazotroph *Herbaspirillum seropedicae* SmR1. Microbiology (N Y). 2013;159(Pt_1):167-175. doi:10.1099/mic.0.061135-0

41. Marin MA, de la Torre J, Ricardo Marques Oliveira A, et al. Genetic and functional characterization of a novel meta-pathway for degradation of naringenin in *Herbaspirillum seropedicae* SmR1. Environ Microbiol. 2016;18(12):4653–4661. doi:10.1111/1462-2920.13313

42. Braune A, Gütschow M, Blaut M. An NADH-Dependent Reductase from *Eubacterium ramulus* Catalyzes the Stereospecific Heteroring Cleavage of Flavanones and Flavanonols. Appl Environ Microbiol. 2019;85(19):e01233–19. doi:10.1128/AEM.01233-19

43. He L, Zhang Z, Lu L, et al. Rapid identification and quantitative analysis of the chemical constituents in Scutellaria indica L. by UHPLC–QTOF–MS and UHPLC–MS/MS. J Pharm Biomed Anal. 2016;117:125–139. doi:10.1016/j.jpba.2015.08.034

44. Xu J, Yang L, Zhao SJ, Wang ZT, Hu ZB. An efficient way from naringenin to carthamidine and isocarthamidine by *Aspergillus niger*. World J Microbiol Biotechnol. 2012;28(4):1803–1806. doi:10.1007/s11274-011-0934-9

45. Fu Q, Tong C, Guo Y, et al. Flavonoid aglycone–oriented data-mining in high-performance liquid chromatography–quadrupole time-of-flight tandem mass spectrometry: efficient and targeted profiling of flavonoids in *Scutellaria barbata*. Anal Bioanal Chem. 2020;412(2):321–333. doi:10.1007/s00216-019-02238-7

46. van den Berg MA, Albang R, Albermann K, et al. Genome sequencing and analysis of the filamentous fungus *Penicillium chrysogenum*. Nat Biotechnol. 2008;26(10):1161–1168. doi:10.1038/nbt.1498

47. Frydoonfar HR, McGrath DR, Spigelman AD. The variable effect on proliferation of a colon cancer cell line by the citrus fruit flavonoid Naringenin. Colorectal Disease. 2003;5(2):149–152. doi:10.1046/j.1463-1318.2003.00444.x

48. Harmon AW, Patel YM. Naringenin inhibits glucose uptake in MCF-7 breast cancer cells: A mechanism for impaired cellular proliferation. Breast Cancer Res Treat. 2004;85(2):103–110. doi:10.1023/B:BREA.0000025397.56192.e2

49. Dunstan MS, Robinson CJ, Jervis AJ, et al. Engineering *Escherichia coli* towards de novo production of gatekeeper (2S)-flavanones: naringenin, pinocembrin, eriodictyol and homoeriodictyol. Synth Biol. 2020;5(1):ysaa012. doi:10.1093/synbio/ysaa012

50. Sordon S, Madej A, Popłoński J, et al. Regioselective ortho-Hydroxylations of Flavonoids by Yeast. J Agric Food Chem. 2016;64(27):5525–5530. doi:10.1021/acs.jafc.6b02210

51. Gao S, Zhou H, Zhou J, Chen J. Promoter-Library-Based Pathway Optimization for Efficient (2 *S*)-Naringenin Production from *p* −Coumaric Acid in *Saccharomyces cerevisiae*. J Agric Food Chem. 2020;68(25):6884–6891. doi:10.1021/acs.jafc.0c01130

52. Gao S, Lyu Y, Zeng W, Du G, Zhou J, Chen J. Efficient Biosynthesis of (2 *S*)-Naringenin from *p* - Coumaric Acid in *Saccharomyces cerevisiae*. J Agric Food Chem. 2020;68(4):1015–1021. doi:10.1021/acs.jafc.9b05218

53. Cui H, Song MC, Lee JY, Yoon YJ. Microbial production of O-methylated flavanones from methylated phenylpropanoic acids in engineered *Escherichia coli*. J Ind Microbiol Biotechnol. 2019;46(12):1707–1713. doi:10.1007/s10295-019-02239-6

54. Leonard E, Yan Y, Fowler ZL, et al. Strain Improvement of Recombinant *Escherichia coli* for Efficient Production of Plant Flavonoids. Mol Pharm. 2008;5(2):257–265. doi:10.1021/mp7001472

55. Palmer CM, Miller KK, Nguyen A, Alper HS. Engineering 4-coumaroyl-CoA derived polyketide production in *Yarrowia lipolytica* through a β-oxidation mediated strategy. Metab Eng. 2020;57:174–181. doi:10.1016/j.ymben.2019.11.006

56. Behzad S, Sureda A, Barreca D, Nabavi SF, Rastrelli L, Nabavi SM. Health effects of phloretin: from chemistry to medicine. Phytochemistry Reviews. 2017;16(3):527–533. doi:10.1007/s11101-017-9500-x

57. Mouradov A, Spangenberg G. Flavonoids: a metabolic network mediating plants adaptation to their real estate. Front Plant Sci. 2014;5:620. doi:10.3389/fpls.2014.00620

