## Supplementary Figures and Tables for "Heterologous naringenin production in the filamentous fungus *Penicillium rubens*"

### Supplementary materials

#### *Sequences of synthetic genes*

Pc4CL *(Petroselinum crispum*, GenBank accession number KX671122.1):

ATGGGTGACTGCGTTGCCCCGAAAGAGGATCTGATCTTCCGCAGCAAACTGCCGGACATTTACATTCCAAAGCATCTGCCGCT
GCATACCTATTGTTTTGAGAACATCAGCAAGGTTGGCGACAAGAGCTGTCTGATCAACGGCGCAACCGGCGAAACCTTTACCT
ACAGCCAGGTTGAGCTGCTGTCCCGTAAAGTTGCCAGCGGCCTGAACAAGCTGGGCATTCAACAAGGTGATACCATTATGCTG
CTGCTGCCGAACTCCCCGGAGTACTTTTTCGCTTTCCTGGGTGCGAGCTATCGCGGTGCAATCAGCACTATGGCGAACCCATT
CTTTACCAGCGCAGAAGTGATCAAGCAACTGAAAGCGAGCCAAGCGAAGCTGATTATCACCCAGGCATGCTATGTTGACAAGG
TTAAGGACTACGCAGCGGAGAAAAACATCCAGATCATTTGTATTGACGATGCACCGCAGGATTGCCTGCACTTTAGCAAGCTG
ATGGAAGCGGATGAGAGCGAAATGCCGGAAGTGGTTATTAACAGCGATGATGTGGTGGCACTGCCGTACAGCTCTGGCACCAC
CGGCCTGCCGAAAGGCGTTATGCTGACCCACAAGGGTCTGGTTACCAGCGTTGCACAACAGGTGGATGGTGATAACCCGAACC
TGTATATGCACTCCGAGGATGTTATGATCTGCATCCTGCCACTGTTCCATATCTATAGCCTGAACGCTGTTCTGTGTTGTGGT
CTGCGTGCGGGCGTTACCATTCTGATCATGCAAAAGTTCGACATTGTGCCGTTTCTGGAGCTGATTCAGAAGTATAAGGTTAC
CATTGGTCCGTTTGTTCCGCCGATCGTGCTGGCCATCGCGAAAAGCCCGGTTGTTGACAAGTACGACCTGTCTAGCGTGCGCA
CCGTTATGAGCGGTGCAGCGCCGCTGGGTAAAGAGCTGGAGGACGCTGTTCGTGCGAAATTCCCGAACGCGAAGCTGGGTCAA
GGCTATGGCATGACCGAAGCCGGTCCGGTTCTGGCGATGTGTCTGGCGTTCGCCAAAGAGCCGTATGAGATTAAGTCTGGCGC
ATGCGGTACCGTTGTGCGTAACGCCGAGATGAAAATCGTTGACCCAGAAACCAACGCGTCTCTGCCGCGTAACCAGCGTGGTG
AGATTTGCATCCGTGGTGATCAGATTATGAAAGGTTACCTGAACGACCCGGAAAGCACCCGCACCACCATCGACGAAGAGGGT
TGGCTGCACACCGGTGACATTGGTTTCATCGACGATGACGATGAACTGTTCATTGTTGATCGTCTGAAAGAAATCATTAAGTA
CAAAGGTTTTCAAGTTGCTCCGGCGGAGCTGGAAGCACTGCTGCTGACCCACCCGACCATCAGCGATGCCGCGGTGGTTCCGA
TGATTGACGAGAAAGCGGGTGAAGTGCCAGTGGCGTTTGTTGTGCGTACCAACGGTTTTACCACCACCGAAGAAGAAATCAAA
CAATTTGTGAGCAAACAGGTTGTGTTCTACAAACGTATCTTCCGCGTTTTCTTCGTTGACGCTATTCCGAAATCCCCGAGCGG
CAAGATTCTGCGTAAGGATCTGCGCGCTCGTATTGCGAGCGGCGACCTGCCGAAGTAA

PhCHS (*Petunia hybrida,* GenBank accession number KP284563.1): codon optimization for *E. coli*

ATGGTGACCGTGGAAGAATACCGTAAGGCGCAACGTGCGGAAGGCCCGGCGACCGTGATGGCGATTGGCACCGCGACCCCGAG
CAACTGCGTTGACCAGAGCACCTACCCGGATTTCTATTTTCGTATTACCAACAGCGAGCACAAAACCGACCTGAAGGAAAAAT
TCAAGCGTATGTGCGAGAAGAGCATGATTAAGAAACGTTACATGCACCTGACCGAGGAAATCCTGAAAGAGAACCCGAGCATG
TGCGAATATATGGCGCCGAGCCTGGACGCGCGTCAGGATATCGTGGTTGTGGAAGTGCCGAAACTGGGCAAAGAGGCGGCGCA
GAAAGCGATTAAGGAATGGGGTCAACCGAAAAGCAAGATCACCCACCTGGTTTTCTGCACCACCAGCGGCGTGGACATGCCGG
GTTGCGATTACCAACTGACCAAACTGCTGGGCCTGCGTCCGAGCGTTAAGCGTCTGATGATGTATCAGCAAGGTTGCTTTGCG
GGTGGCACCGTGCTGCGTCTGGCGAAAGATCTGGCGGAAAACAACAAGGGTGCGCGTGTTCTGGTTGTGTGCAGCGAGATTAC
CGCGGTGACCTTCCGTGGCCCGAACGACACCCACCTGGATAGCCTGGTTGGTCAGGCGCTGTTTGGTGATGGTGCGGGTGCGA
TCATTATCGGCAGCGATCCGATTCCGGGTGTTGAGCGTCCGCTGTTCGAACTGGTGAGCGCGGCGCAAACCCTGCTGCCGGAC
AGCCATGGTGCGATTGATGGTCACCTGCGTGAAGTTGGCCTGACCTTTCACCTGCTGAAAGACGTGCCGGGTCTGATTAGCAA
AAACATCGAGAAGAGCCTGGAGGAAGCGTTCAAGCCGCTGGGCATTAGCGACTGGAACAGCCTGTTTTGGATTGCGCACCCGG
GTGGCCCGGCGATTCTGGATCAAGTTGAAATCAAACTGGGCCTGAAGCCGGAGAAACTGAAGGCGACCCGTAACGTTCTGAGC
AACTACGGTAACATGAGCAGCGCGTGCGTGCTGTTTATCCTGGATGAAATGCGTAAAGCGAGCGCGAAAGAGGGTCTGGGTAC
CACCGGCGAGGGTCTGGAATGGGGTGTGCTGTTCGGCTTTGGTCCGGGCCTGACCGTGGAAACCGTTGTTCTGCATAGCGTTG
CGACCTAA

#### Supporting Tables

**Table S1.** Sequences of primers used in the study.

| Primer | Sequence (5' to 3') | Application |
|---|---|---|
| lvl0_Pc4CL_F | TTGAAGACTTAATGGGTGACTGCGTTGCC | Building of pFL_0_1_Pc4CL |
| lvl0_Pc4CL_R | AAGAAGACAAAAGCTTACTTCGGCAGGTCGCCG | |
| lvl0_PhCHS_F | TTGAAGACTTAATGGTGACCGTGGAAGAATACCG | Building of pFL_0_2_PhCHS |
| lvl0_PhCHS_R | AAGAAGACAAAAGCTTAGGTCGCAACGCTATGCAGAAC | |
| lvl0_scr_F | AGTCAGTGAGCGAGGAAGC | Colony PCR for level 0 vectors |
| lvl0_scr_R | AATAGGCGTATCACGAGGC | |
| lvl1_scr_F | CACATTGCGGACGTTTTTAATGTACTG | Colony PCR for level 1 vectors |
| lvl1_scr_R | CCGCCAATATATCCTGTCAAACACTG | |
| Pc4CL_F | TGAATAGAAGACTCGGTGATGCAGCAAATAGCGACTGTTCGTTGCGGGGTCCGAACCCGCTCGGCAGCACCGGGCTCTCCCTACTATCCCTCGATAGCAGCTGCATTGGTCTGCCATTG | Amplification of donor DNA for in vivo HR of Pc4CL_ ergA_ PhCHS into *pen* locus |
| Pc4CL_R | CGCAGGGTTTGAGAACTCCGATCTTAAATCCAAGG | |
| ergA_F | GCAAGGTGCTATTCTAGGTAGGGTATGCCTAGCAATGCCATGATCTTATGACCTCAGTAATCTGACCTTGGATTTAAGATCGGAGTTCTCAAACCCTGCGTGCCTACCGCTCGTACCATGGG | |
| ergA_R | GACCGTCTATAACTCTGGATCCCCCGGGCTGCAG | |
| PhCHS_F | CTCTGCGTCCGTCCGTCTCTCCGCATGCCAGAAAGAGTCACCGGTCACTGTACAGAGCTCGAATTCCTGCAGCCCGGGGGATCCAGAGTTATAGACGGTCCGGCATAGGTAAGGAGAG | |
| PhCHS_R | GAATGCCGATTTGTCGGCAACGAGAGGTATGTCTAAGGTCTCGAGTTTAATTAGAATATTTACTAACAGCGTTTAGGAGCTCTTCCCCTAGCCAACTAGGTGCTTGGGATGTTCCATGG | |
| ergA_F2 | TAGAAGACTCGGTGATGCAGCAAATAGCGACTGTTCGTTGCGGGGTCCGAACCCGCTCGGCAGCACCGGGCTCTCCCTACTATCCCTCGATAGCATACCGCTCGTACCATGGGTTG | Amplification of donor DNA for in vivo HR of ergA_ PhCHS into *pen* locus |
| sgRNA_F | AAAAAAGCACCGACTCGGTGCCACTTTTTCAAGTTGATAACGAACTAGTCTTATTTCAACTTGCTATGCTGTTTCCAGCATAGCTCTGAAAC | sgRNA targed on *pen* locus |
| sgRNA_R | ATGTAATACGACTCACTATA**GAACCAACATCATTAAGCAG**GTTTCAGAGCTATGCTGGAAA | |
| cPen_F | TGTTCGTAGAAGATCTGCCA | Colony PCR of Pc4CL integration |
| c4CL_R | CGCTTGGCTCGCTTTCAGTTGCTTGAT | |
| c4CL_F | ATCGACGATGACGATGAACTGT | Colony PCR of Pc4CL_ ergA_ PhCHS integration |
| cCHS_R | CCTCCAGGCTCTTCTCGATG | |
| cP40s_F | GAGTTATAGACGGTCCGGCATAGGTAAG | Colony PCR of PhCHS integration for 3F |
| cPen_R | ATTGCCAGTCTCACTATCCGATATGC | |
| cP40s_R | CCTTACCTATGCCGGACCGT | Colony PCR of ergA_PhCHS integration for 2F |

Underlined bases: restriction site sequence; Bold: 20 bp target sequence of sgRNA; HR: homologous recombination.

#### Supporting Figures

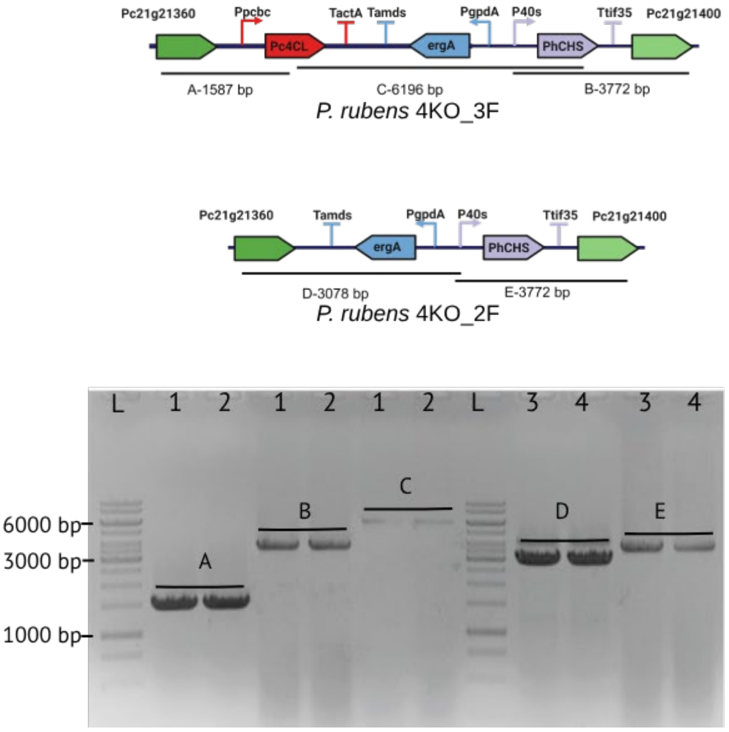

**Figure S1.** Colony PCR verification of integration of naringenin biosynthesis cluster into *P. rubens* 4xKO. Two colonies of the 3F and 2F variants were randomly selected and analyzed for the correct integration using the following PCR primer pairs: A) primers cPen_F and c4CL_R for the 3F variant; B) primers c4CL_F and cCHS_R for the 3F variant; C) primers cP40s_F and cPen_R for the 3F variant; D) primers cPen_F and cP40s_R for the 2F variant; and E) primers cPen_F and c4CL_R for the 2F variant. PCR amplified regions and expected size of amplicons are depicted on top. L: 1kb DNA ladder; 1, 2 are the PCR products of two 3F clones; 3, 4 are the PCR products of two 2F clones.

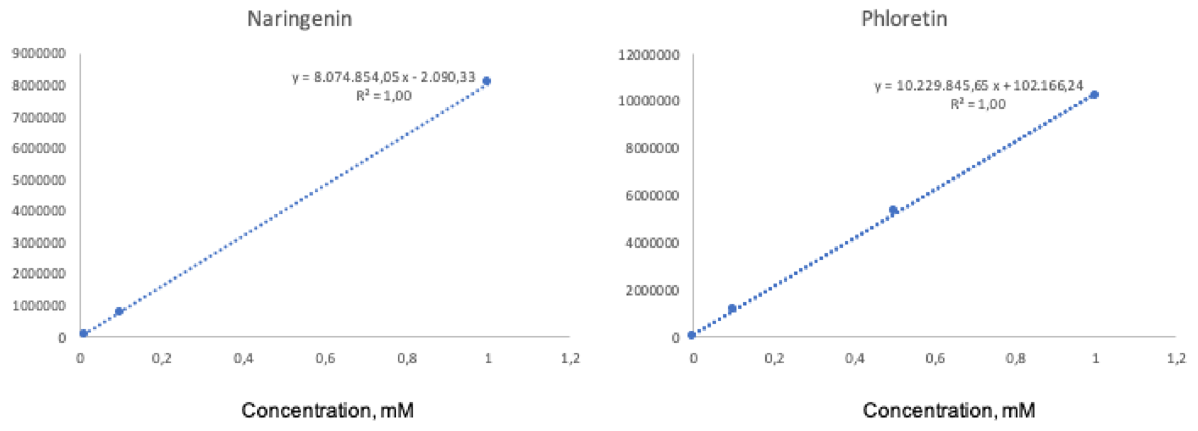

**Figure S2.** Calibration plot of naringenin and phloretin dissolved in DMSO and analyzed by HPLC. The compounds were detected at 288 nm and the analysis was performed as described in the method section. The range of calibration curve was 0.01 mM to 1 mM.

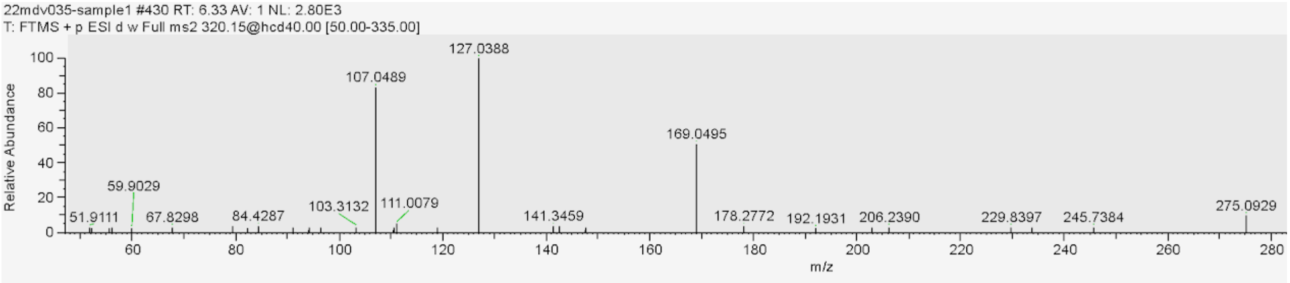

**Figure S3.** High resolution tandem MS of phloretin produced by *P. rubens* 4xKO 3F (275.0929 m/z $[M+H]^+$, RT=6.33).

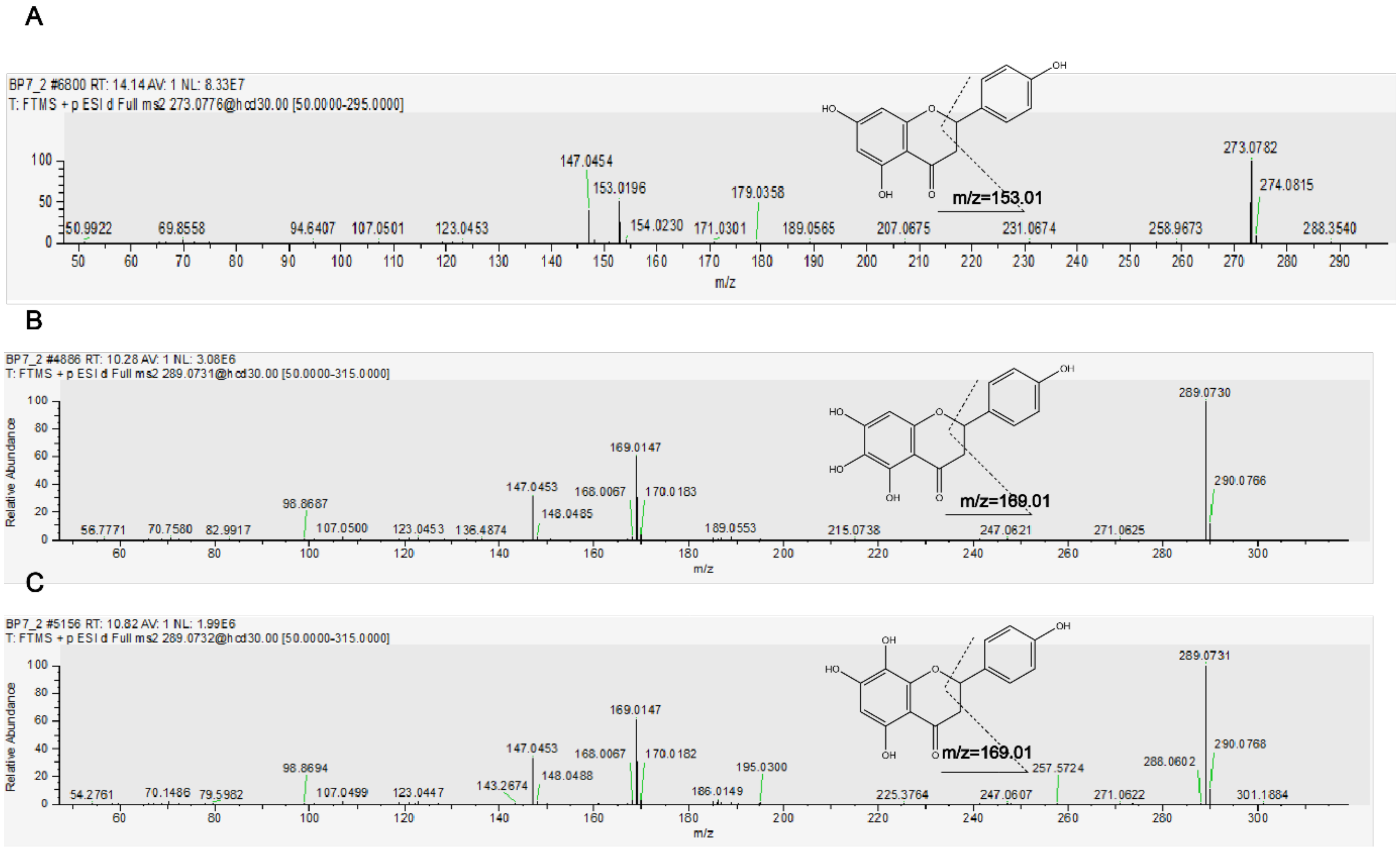

**Figure S4.** Product-ion mass spectra of: naringenin (A) and carthamidin (B), and isocarthamidin (C).
